# Identification of Ovarian Cancer Gene Expression Patterns Associated with Disease Progression and Mortality

**DOI:** 10.1101/473165

**Authors:** Md. Ali Hossain, Sheikh Muhammad Saiful Islam, Julian Quinn, Fazlul Huq, Mohammad Ali Moni

**Author notes:** **Correspondence**: Dept of CSE, Jahangirnagar University, Dhaka, Bangladesh, The University of Sydney, Sydney Medical School, School of Medical Science, Discipline of Biomedical Sciences, NSW 1825, Australia. Funding information.

## Abstract

Ovarian cancer (OC) is a common cause of death from cancer among women worldwide, so there is a pressing need to identify factors influencing mortality. Much OC patient clinical data is now publically accessible (including patient age, cancer site stage and subtype), as are large datasets of OC gene transcription profiles. These have enabled studies correlating OC patient survival with clinical variables and with gene expression but it is not well understood how these two aspects interact to influence mortality. To study this we integrated clinical and tissue transcriptome data from the same patients available from the Broad Institute Cancer Genome Atlas (TCGA) portal. We investigated OC mRNA expression levels (relative to normal patient tissue) of 26 genes already strongly implicated in OC, assessed how their expression in OC tissue predicts patient survival then employed Cox Proportional Hazard regression models to analyse both clinical factors and transcriptomic information to determine relative risk of death associated with each factor. Multivariate analysis of combined data (clinical and gene mRNA expression) found age, ovary tumour site and cancer stage IB significantly correlated with patient survival. Univariate analysis also confirmed significant differences in patient survival time when altered transcription levels of KLK6, CD36, MEF2C and SCGB2A1 were evident, while multivariate analysis that considered the 26 genes simultaneously revealed a significant relationship of mortality with KLK6, CD36 and E2F1 genes. However, analysis that considered all 26 genes with clinical variables together identified WFDC2, E2F1, BRCA1, KLK6, SCGB2A1 and SLPI genes as independently related to mortality in OC. This indicated that the latter genes affect OC patient survival, i.e., provided mechanistic and predictive information in addition to that of the clinical traits and provide strong evidence that these genes are critical markers of processes that underlie OC progression and mortality.

## 1 INTRODUCTION

Ovarian carcinoma (OC) accounts for the great majority of cases of cancer affecting the ovaries. It is the fifth most common cause of cancer deaths in women with an estimated 239,000 new cases and 152,000 deaths worldwide annually (Ferlay et al. (2014)) and remains the leading cause of death from gynecologic malignancy (Seigel et al. (2014); Gov and Arga (2017)) with five year mortality rate in the United States of 35% (Gov and Arga (2017)) in 14,000 annual deaths (acs (2018)). Current standard treatment mainly comprises platinum-based chemotherapeutics, however the most prevalent OC type, high grade serous carcinoma (HGSC), often shows high chemotherapy resistance as well as greatly altered genome and transcriptome. High throughput gene and gene expression data from patient tissue is emerging as an important research tool to identify key intrinsic factors that affect cancer behaviour with numerous genetic mutations shown to be associated with OC development and progression. Such genes are of wide interest for their potential prognostic power and as possible drug targets.

While tumour-associated coding gene mutations can affect both gene product function and levels, it is their gene transcript levels that give more direct information regarding levels of their respective protein gene products. The latter is important in determining invasiveness, metastasis establishment, immune evasion and growth at metastatic sites, as well as indirect pathological effects (via secreted factors) on other tissues that impair health. Clearly, OC gene expression patterns will influence patient fate and their quantification may also yield clues to important cellular pathways underlying disease processes. Tumour tissue transcript levels determined by RNA sequencing (RNAseq) are thus becoming commonly used to study tumour clinical features (Moni and Liò (2014)).

Earlier work on OC has focused on influence of clinical features of the patient on outcomes (Zheng et al. (2009); Brown and Frumovitz (2014); Moni et al. (2014); Mascarenhas et al. (2006); Zhang (2016); Moni and Lio (2014)), which have identified significant factors such as age at diagnosis, age at menopause, disease stage and histological grade, tumour size, type of therapy received and family history of OC. In addition, new transcriptomic data from tumour tissue has identified a number of potential gene expression markers for patient progression (Zhang et al. (2013); Lynch et al. (2009); Moni and Lio’ (2017); Friedenson (2005); Tecza et al. (2015); Moni and Lio (2015)). However, generally less studied until recently (Ahmed and Begum (2018)) is how such tumour transcript levels may interact with clinical factors to affect cancer outcomes, which is a crucial consideration in determining how to use and interpret the transcript data. Publicly available tumour transcript datasets are increasing in number along with clinical datasets, and this makes feasible large scale investigations of interaction between transcript levels and clinical trait. In particular, studies on combined datasets (containing both types of data from the same patient) can be particularly powerful tools for revealing important features of gene function in the tumour context, and informing the clinical use of such data. We thus employed the rich database available from The Cancer Genome Atlas (TCGA) project, a collaboration between the National Cancer Institute (NCI) and National Human Genome Research Institute (NHGRI), and supported by the Broad Institute in order to make available the survival, clinical, and gene expression data to analyse these either separately or in combination, for the purpose of clarifying which type of mutual influences are important to OC patients survival as well as which factors are potent independent predictors. We selected 26 genes of interest previously validated as important in OC or OC models.

The influence of both clinical factors and disease marker gene expression on ovarian cancer patient survival can now be determined using standard Cox Proportional Hazard (PH) (Cox (1992)) models for univariate analysis, multivariate analysis (Gabriel and Glavin (1978)) and the penalized Cox PH model (Heinze and Schemper (2001)) for combined analysis (Ahmed and Begum (2018)). Important clinical features with associations to the disease can be identified and selected by consulting the OC literature for use in these models. In this study, Cox PH regression modelling is employed for the analysis or post-diagnosis survival time. Clinical variables that were selected for analysis included age, ethnicity, anatomical site of cancer, histological grade of cancer, primary tumour site and neoplasm status with tumour were considered from the literature of OC. We focused on research work that investigated associations between gene expression, clinical factors and survival in patients with OC. We performed univariate and multivariate analysis for selected significant 26 genes selection through a process of literature search and evaluation, and we also ran combined analysis for the 26 genes and clinical factors on OC patient survival. We gathered survival, clinical, and gene expression data from TCGA separately and combined them in order to assess the joint role of genetic and clinical factors.

Our analysis followed the methodology of Xu and Moni (2015) (Xu et al. (2015)) who used Cox PH regression modelling for the analysis of post-diagnosis survival time. We performed an analysis to identify most significant genes among selected 26 genes as well as clinical variables, affecting patients survival in OC. To do so, we used product-limit estimator to estimate survival function for each gene separately, then used log rank test to detect which genes differs in expression levels significantly between altered and not altered group. Finally we performed univariate Cox hazard regression analysis to determine each gene’s likelihood of contribution to deaths and two multivariate Cox PH regression analyses, the first taking consideration of 26 genes simultaneously and with the second considering all 26 genes and clinical variables to determine most significant genes and clinical variables in context of likelihood of risk of death.

## 2 METHODS & MATERIALS

### Data

We collected the RNAseq data for this study from TCGA genome data analysis centre (http://gdac.broadinstitute.org/) which is an interactive data system for researchers to search, upload, download, and analyse cancer genomic data (Tomczak et al. (2015)). Since our goal was to explore a particular point of interest, survival analysis of OC on clinical and genetic factors, we retrieved the anonymized clinical data and RNAseq data for OC (Ovarian Serous Cystadeno Carcinoma TCGA, Provisional) from the cBioPortal (Cerami et al. (2012)).

In Clinical dataset there are 577 cases with 87 features. Cases that had RNAseq gene expression data included 535 cases with 5689 genes. We employed six clinical factors (ethnicity, anatomical site of cancer, histological grade of cancer, primary tumour site, and neoplasm status with tumour) along with 26 genes commonly cited as significant factors in the OC literature (Table 1) (Adib et al. (2004); McLemore et al. (2009); SJ. (2017); tar (2014)). Using these data we investigated a single outcome variable, namely OC-specific survival. 48 OC patient records that did not include clinical information were excluded from this analysis. We matched patient ID in both clinical and RNAseq dataset and identified 529 patients with data available for both. Among the clinical variables six clinical variables given above were considered (Moorman et al. (2009); McLemore et al. (2009)). Tumour histological subtype and age at initial diagnosis were collected from pathology reporting. OC stage was recorded according to American Joint Committee on Cancer (AJCC) staging classification (acs (2017)). Patient survival data was taken from overall number of months of patient survival but converted to days of survival. Normal and tumour samples were identified using the TCGA barcode; two digits at position 14-15 of the barcode denote the sample type. Tumour type spans from 01 to 09, normal type from 10 to 19 and control samples from 20 to 29. Fold change (FC) was used as a common approach in differential gene expression analysis between two conditions (?). For example, if gene expression read counts (determined by RNAseq) for a patient was 60 while that of a normal patient was 30, patient gene FC was 2. FC is most suitable when the gene expression distribution is symmetric. However, in RNAseq analyses, expression levels are modelled by the discrete counts of reads mapped to a known gene in a reference genome. Poisson and Negative binomial distribution assumptions are taken for reading counts. When the abundance rate of a particular gene expression level is very low, read count distributions modelled by the Poisson or Negative binomial are skewed to the right. Thus, as using FC as a measure of differential expression may not be appropriate in such cases we transformed the gene expression value using a standardizing transformation and calculated z-scores for each expression value. For the expression from RNAseq experiments, the standard rule is to compute the relative expression of an individual gene in tumour samples using gene expression distribution in a reference population. A reference population was considered either all diploid tumours for the gene in question or, when available, normal adjacent tissue. The resulted value (z-score) indicates the number of standard deviations away from the mean expression in the reference population. Here, we considered genes showing *z* > ±1: 96 to be differentially expressed. We computed z-scores for RNAseq data using following formula:

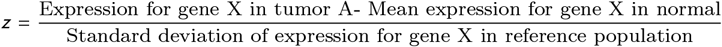

**TABLE 1.** Ovarian cancer associated significant 26 genes and their sources.

| Gene Name | Resource | Reference |
| --- | --- | --- |
| BRCA1, BRCA2, BSCL2, CTNNB1, ERBB2, MSH2, MUC16, PIK3CA, KRAS, TP53 | http://www.cancer-genetics.org/X1003.htm | SJ. (2017) |
| KRAS, PIK3CA, and TP53 | Mutation profile and clinical outcome of mixed endometrioid-serous endometrial carcinomas are different from that of pure endometrioid or serous carcinomas | Coenegrach et al. (2015) |
| KLK6, CP, SLPI, EZH2, HPN, SCGB2A1 | Predicting biomarkers for ovarian cancer using gene-expression microarrays | Adib et al. (2004) |
| TLR4 | The inflammatory microenvironment in epithelial ovarian cancer: a role for TLR4 and MyD88 and related proteins | Li et al. (2016) |
| WFDC2 | HE4 (WFDC2) gene overexpression promotes ovarian tumor growth | Moore et al. (2014) |
| RXRG, DVL1, MEF2B, MEF2C, BSCL2, CD36, RXRG, E2F1 and TLR4 | The impact on high-grade serous ovarian cancer of obesity and lipid metabolism-related gene expression patterns: the underestimated driving force affecting prognosis | Cuello et al. (2018) |

We thus used the z-score values to define samples with “altered” and “normal” (unaltered) expression of a gene. We have assumed a sample to be altered if the z-score for that sample is equal to or higher than a specific threshold value such as z=2, as noted. We therefore define altered versus normal as follows:

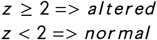

### Methods

We performed the following analysis related to survival of patients with OC. First we used product-limit estimator for estimation of survival function, then we performed log rank test to determine whether the survival functions of two different groups (patients with altered gene expression and patients with unaltered gene expression) exhibit statistically significant difference and after then we used Cox Proportional Hazards (Cox PH) regression models to determine the significance of genes, clinical factors on comparative risk of death and finally we performed functional analyses of our most significant genes found from our analysis. We selected important clinical and demographic variables affecting OC by literature review and similarly selected 26 genes having experimentally determined significance in context of OC. The expression z-score for each gene was identified as being in “altered” or “normal” categories based on a significance threshold value (*z* < 2) as noted in the Data section above. We performed Cox PH regression for every gene individually known as univariate regression as well as multivariate analysis taking all 26 genes simultaneously and finally multivariate regression on combined set of clinical and gene expression data. Survival analysis is a statistical analysis for estimating expected duration of time until one or more events happen, such as death in cancer and failure in mechanical systems. Normally, survival analysis is carried in three steps: determining time to event, formulating a censoring indicator for subject inclusion in the analysis, and the time to occurrence of event. Censoring in survival analysis is usually done in two ways, right censoring and left censoring. Right censoring occurs when a subject leaves the study before an event occurs, or the study ends before the event has occurred. Left censoring happens when the event of interest has already occurred before enrollment. This is very rarely encountered. Right censoring is again of two types. First one is Type I right censoring results from completely random dropout (e.g emigration) and/or end of study with no event occurrence and the second one, Type II right censoring, occurs with end of study after fixed number of events amongst the subjects has occurred.

In survival analysis, one is interested in estimation of survival function of samples divided into subgroups or as whole. There are two types of estimation of it, one is non parametric in which no prior assumption of distribution of survival time is made and other is parametric estimation in which there are some assumption along with assuming predefined distribution of survival time. As stated above, we used non parametric technique for estimation of survival function. There are several such techniques, we used product-limit estimator. In short product-limit (PL) estimator of the survival function is defined as follows:

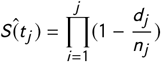

Here 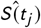 is estimated survival function at time *t_j_*, *d_j_* is the number of events occurred at *t_j_*, and *n_j_* is the number of subjects available at *t_j_*. After estimating survival function, two or more groups can be compared using log-rank test. For example, we used Log-rank test to detect the most significant genes in the case of patient’s survival time in altered versus unaltered groups in context of gene expression. The null hypothesis is following: *H*_0_: Survival function for patients with altered gene expression is not different from the patients with normal (unaltered) gene expression

*H_A_*: Survival functions are different for these two groups. Symbolically these can be written

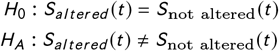

Survival analysis methods can also be extended to assess several risk factors simultaneously similar to multiple linear and multiple logistic regression analysis. One of the most popular regression techniques for survival analysis is Cox PH regression, which is used to relate one or several risk factors or exposures, considered simultaneously, to survival time. In a Cox proportional hazards regression model, the measure of effect is the hazard rate, which is the risk of failure (i.e., the risk or probability of suffering the event of interest), given that the participant has survived up to a specific time. Using Cox PH regression, first we performed univariate survival analysis by selecting each gene separately, then conducted a multivariate survival analysis taking all 26 genes simultaneously and finally fit a Cox PH model on all of selected six clinical factors and 26 genes combined, modelling the hazards of having the event under investigation (Ovarian Serous Cystadeno carcinoma, in this case), using an undetermined baseline hazard function and an exponential form of a set of covariates. Mathematically we can write the model as following:

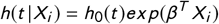

Whereas *h*(*t|X*) is the hazard function conditioned on a subject i with covariate information given as the vector *x_i_*, *h*_0_(*t*) is the baseline hazard function which is independent of covariate information, and represents vector of regression coefficients to the covariates correspondingly. We have calculated the hazard ratio (HR) based on the estimated regression coefficients from the fitted Cox PH model to determine whether a specific covariate affects patient survival. The hazard ratio for a covariate *x_r_* can be expressed by the following simple formula exp (*β_r_*). Thus, the hazard ratio for any covariate can be calculated by applying an exponential function to the corresponding (*β_r_*) coefficient.

Pathways and gene ontology (GO) for these genes were analysed using KEGG pathway database (*http://www.genome.jp/kegg/p*) and enriched using (*http://amp.pharm.mssm.edu/Enrichr/enrich*), a web based software tool.

## 3 RESULTS AND DISCUSSION

6 clinical factors (including age) and mRNA expression data for 26 genes were employed in our analysis. All clinical factors were categorical except age. Age distribution of patients is shown in Figure 1. Average age of the patients at the time of their diagnoses was 59.7 years, with a range between 26 and 89 years old. The descriptive summary statistics of these factors shown in Table 2. From Table 2, we observe that the OC stage variable has the highest number (10) of categories and 71.03% of all patients are of stage IIIC. The next largest group is stage IV with 15.36% cases. There are 8 categories for histology type with the highest percentage of patients (84.67%) from was G3, i.e., poorly differentiated, while the second largest group (12.02%) of patients had moderately differentiated grade. Regarding ethnicity, most patients (85.27%) are from the European-caucasian descent population, while African American is next largest group with 5.89%. In the case of OC anatomical site, most (72.53%) women had bilateral cancers, while cancer patients with left and right ovary are 14.65% and 12.82% respectively. Most women had cancer in the ovary itself (99.13%), the remaining minor percentage of women had tumours located in omentum and/or peritoneum.

**FIGURE 1.**
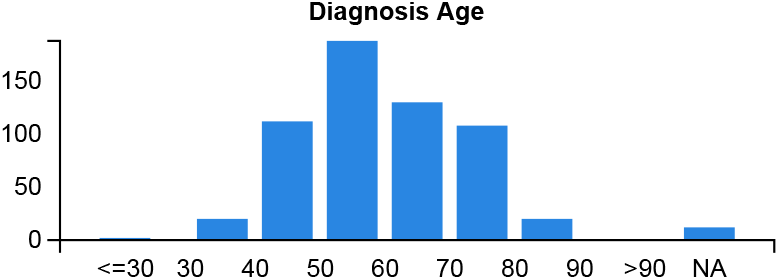
Distribution of Age at Onset of Diagnosis

**TABLE 2.**
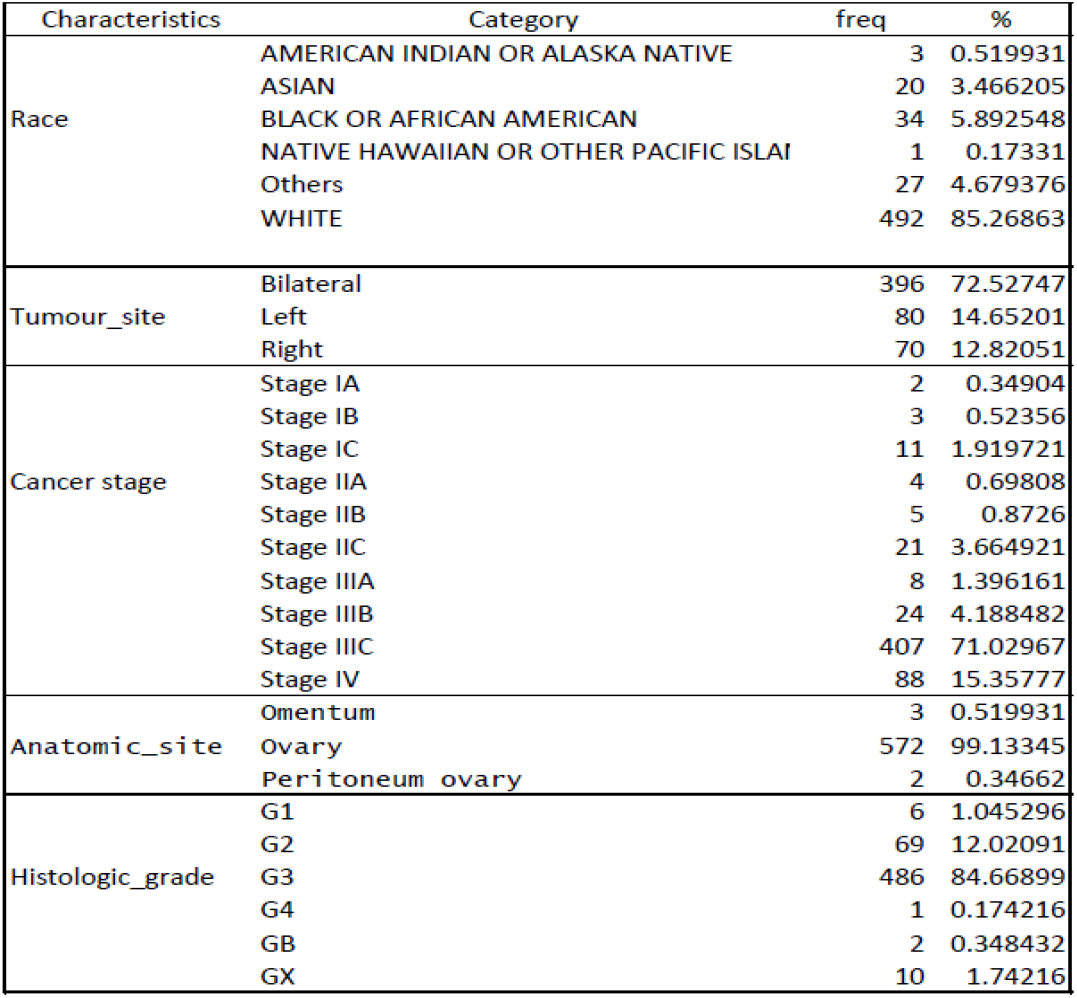
Descriptive Statistics of Clinical Predictors

### 3.1 Survival pattern for gene expression data

We have estimated survival function for altered and unaltered group for each of the 26 genes by applying product limit (PL) estimator. We then compared estimated survival function for altered and unaltered group using log rank test. Those genes for which there is statistically significant difference are shown below. The significant role for these genes is indicated by their p-values in differential survival pattern when comparing their expression level in two categories (altered and unaltered). Surprisingly p53 gene, which is known for its implication in cancer in general, has p-vlaue greater than 0.05. From Figure 2, we can see that patients having altered expression of CD36, MEFC2, KLK6 and SCGB2A1 genes are less likely to survive compared to the non-altered group. Note that, the red line in the graphs indicates normal gene expression and the blue line indicates the altered gene expression group.

**FIGURE 2.**
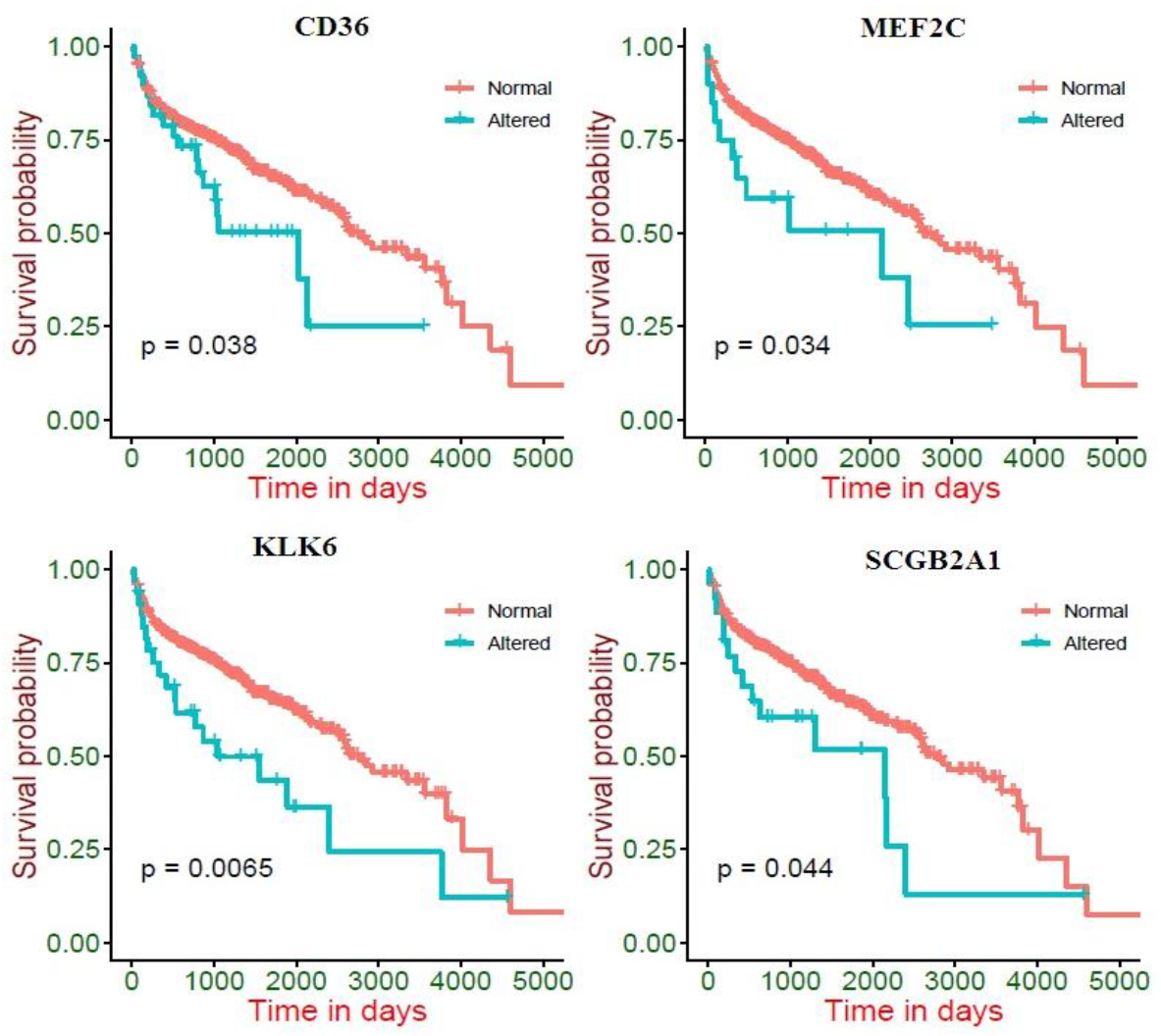
Survival pattern of altered and normal(non-altered) groups for CD36, MEF2C, KLK6, and SCGB2A1

### 3.2 Modeling the hazard risk on the RNA-Seq data

One can measure the relative likelihood of risk of death in OC for each gene separately as well as for all genes simultaneously. Consequently one can determine which genes are most significant in case of survival of patients in OC. For these purposes, Cox PH regression model is used. We considered both univariate (separately for every gene) and multivariate (incorporating rest of the genes) for Cox PH model on each of the 26 genes. Table 3 shows the estimated Coefficients (*β*), with corresponding hazard ratios (HR), and p-values from those analyses. We noted that the p-values for genes CD36 and KLK6 showed that their expression profile have statistically significant association with OC patient survival in both univariate and multivariate analyses. In contrast, genes SCGB2A and MEF2C showed significant association only in univariate analysis, while E2F1 showed significance only in multivariate analysis. Surprisingly, even though the KRAS gene has previously been shown by experimentation to have a significant role in OC (Ratner et al. (2010)), we did not find its significance in our analysis, both in univariate and multivariate analysis.

**TABLE 3.**
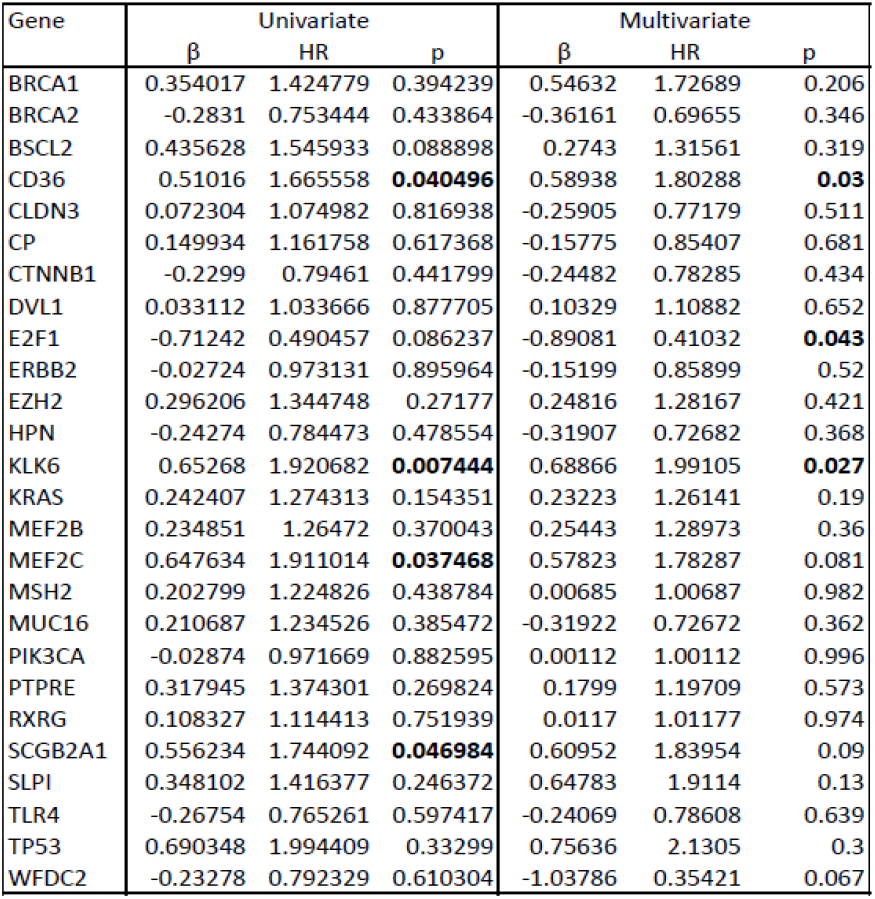
Summary of Univariate and Multivariate Cox Proportional Hazard Model for mRNASeq data.

### 3.3 Modeling hazard on the combined model containing both clinical and RNAseq data

We have performed multivariate Cox PH regression analysis with both clinical and RNA-Seq data simultaneously. We have presented estimated regression coefficients in Table 4 along with hazard ratios (HR),z-values and corresponding p-values in our combined Cox PH model. Here, n= 497, number of events (deaths)= 178 after deleting 32 observations due to missing data values. Table 4 shows the summary of the Cox Proportional Hazard Model result. We observe from Table 4 that if the patient’s age increases one-year, likelihood of hazard increases 1.02 times. Notably, we found no evidence that ethnicity, histologic grade and anatomical OC site have any statistically significant association with hazard. Surprisingly, our analysis indicates that patients with a right side ovary tumour have a higher risk of death (2.27 times) compared to patients with a bilateral tumour, but patients having left side and bilateral tumour exhibit same likelihood of risk of death. Tumour stage is an important factor affecting risk of death in many cancers, and consistent with this, we found that patients with Stage IB OCare 29.5 times more likely to die compared to stage IA, although there was no evidence that other OC stages exerted a significant influence on the likelihood of death compared to stage IA. We then conducted our hazard analysis using the 26 selected genes. Among these, only WFDC2, E2F1, BRCA1, KLK6, SCGB2A1 and SLPI expression levels showed a significant association to patient survival. Patients with altered WFDC2 and E2F1 gene expression have respectively a 0.272 and 0.287 times lower risk of death compared to those with no such alterations. In contrast, patients with altered BRCA1, KLK6, SCGB2A1 and SLPI expression have respectively 2.42, 1.91, 2.48 and 2.62 times higher rates of likelihood of death than those with no alteration in expression in these genes. A Venn diagram of the significant genes summarises the relationships between altered gene expression and risk of death identified using different methodologies and is shown in figure 3. From univariate, multivariate analysis and Cox PH regression analysis on combined data, we found that 8 genes among the 26 studied showed significant association with risk of death; these were KLK6, E2F1, WFDC2, BRCA1, SLPI, SCGB2A1, CD36 and MEF2C. Gene expression of CD36 was significant in both univariate and multivariate analysis, that of E2F1 in multivariate and combined hazard models, and SCGB2A1 expression in both univariate and combined hazard model. Notably, KLK6 gene expression was found significant in all of the three types of models. One gene, MEF2C, was significant only in univariate analysis and expression of three genes (WFDC2, BRCA1, SLPI) were significant in combined hazard model only. The latter suggest that knowledge of these gene expression levels does not add to information from the clinical data.

**TABLE 4.**
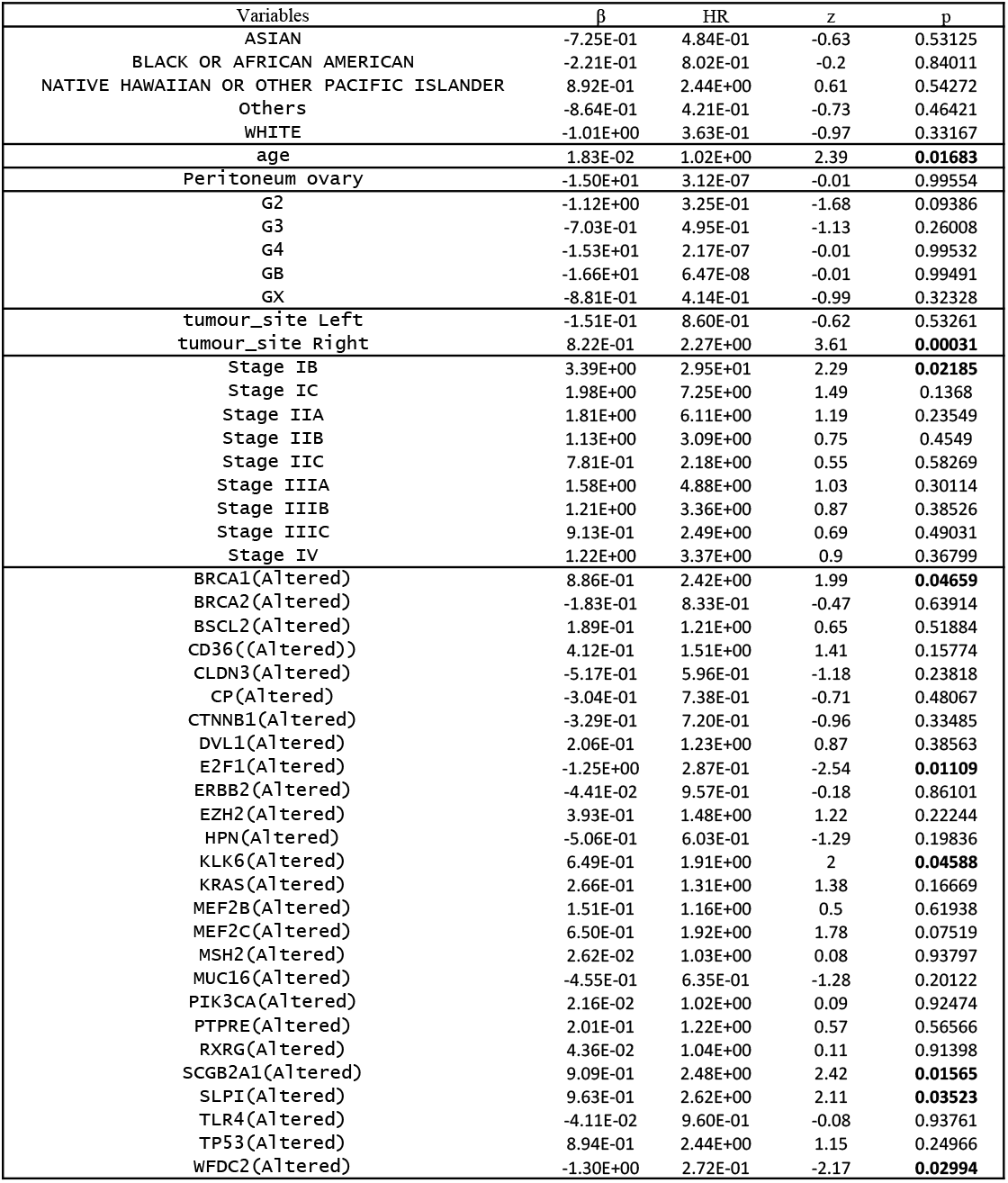
Summary of the Cox Proportional Hazard Model for combined mRNAseq and Clinical data

**FIGURE 3.**
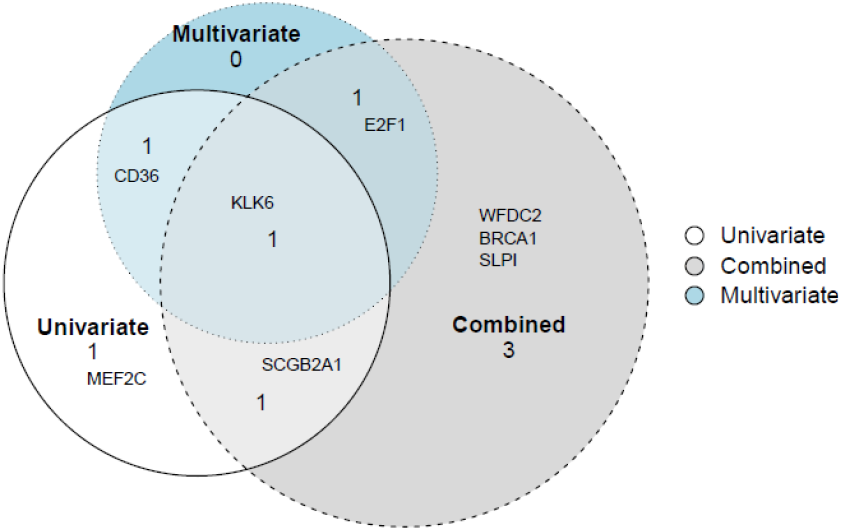
Venn diagram of significant genes found in univariate, multivariate and combined hazad model analysis

**FIGURE 4.**
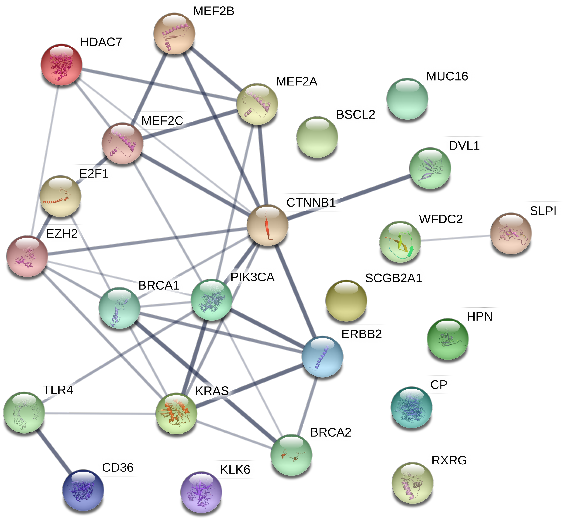
PPI network of 26 selected genes of OC

### 3.4 Pathway and functional correlation analysis of the significant genes

We observed that twenty significant pathways including microRNAs in cancer, bladder cancer, non-small cell lung cancer, and pancreatic cancer are associated with the significantly regulated genes for Ovarian Cancer. Genes associated with these pathways and corresponding p-values are presented in the table (see Table 5).

**TABLE 5.**
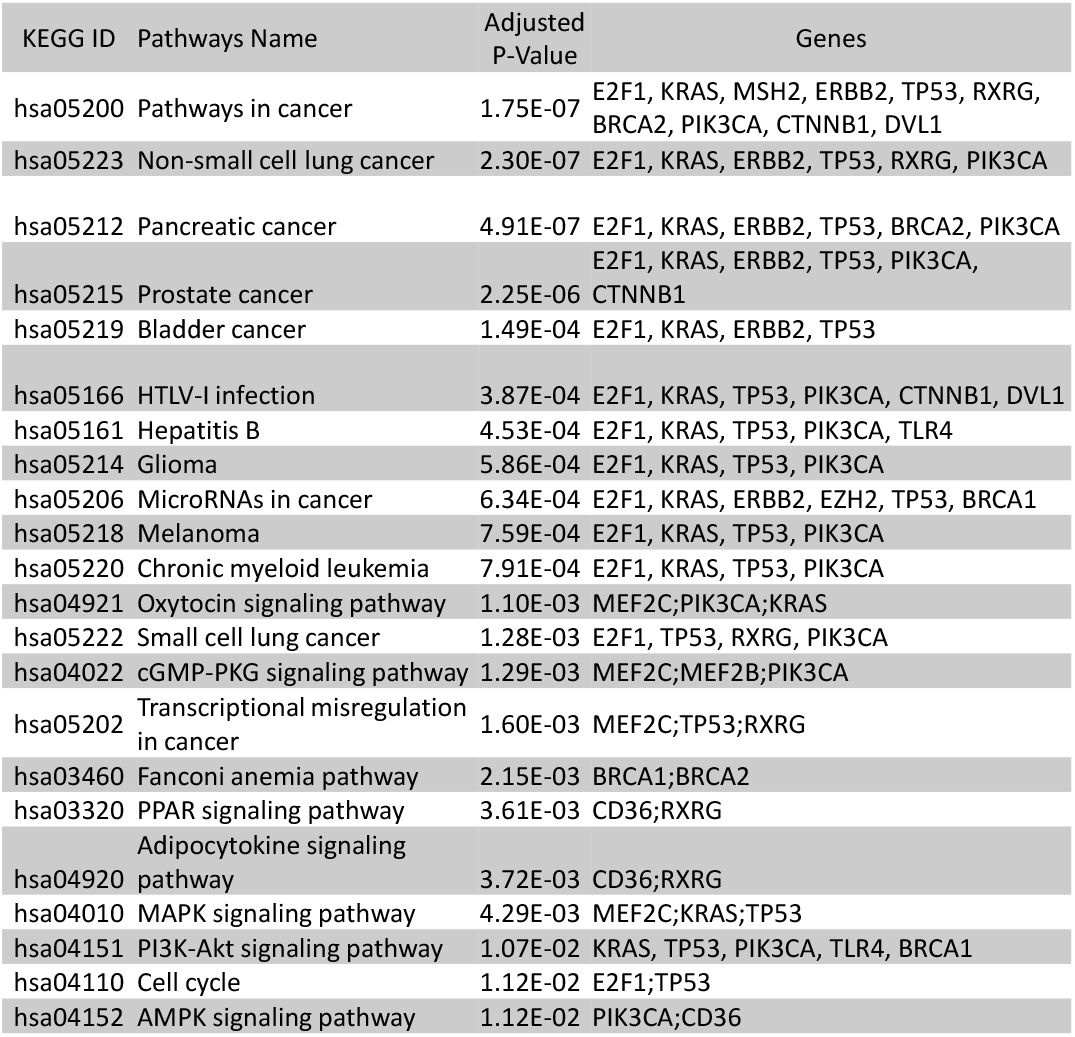
Pathway associated with the selected 8 significantly associated genes with the Ovarian Cancer. Twenty two KEGG pathways were found using Enrichr for these genes. Genes associated with these pathways and corresponding Adjusted p-values are presented in this table.

We also performed the biological process ontology enrichment analysis (see table 6) of these identified significant genes using the Enrichr software tool. We found 18 biological pathways associated with these significant genes as shown in Table 6. We have investigated protein protein interaction network generated using STRING Szklarczyk et al. (2016), a web-based visualization software resource. Most of the key genes are connected with each other through the PPI network. However some genes are not connected with the main network.

**TABLE 6.** Gene Ontolgy terms with underlying Biological pathways associated with 8 significant genes in context of the Ovarian Cancer along. Eighteen Gene Ontology terms along with biological pathways,found significantly associated with these significant genes with adjusted p-values shown in this table.

| GO Term | Pathways | Adjusted P-value | Genes |
| --- | --- | --- | --- |
| GO:0010604 | positive regulation of macromolecule metabolic process | 6.37E-07 | MEF2C;HPN;E2F1;KRAS;CD36;BRCA1;TLR4;TP53 |
| GO:0043406 | positive regulation of MAP kinase activity | 8.97E-05 | MEF2C, KRAS, ERBB2, EZH2 |
| GO:0008630 | intrinsic apoptotic signaling pathway in response to DNA damage | 9.44E-05 | MSH2;E2F1;BRCA1TP53 |
| GO:0006978 | DNA damage response, signal transduction by p53 class mediator | 3.33E-03 | BRCA1;TP53 |
| GO:0097193 | intrinsic apoptotic signaling pathway | 5.09E-03 | E2F1;BRCA1;TP53 |
| GO:0009891 | positive regulation of biosynthetic process | 6.08E-03 | CD36;TLR4 |
| GO:1904645 | response to amyloid-beta | 6.59E-03 | CD36;TLR4 |
| GO:0002755 | MyD88-dependent toll-like receptor signaling pathway | 7.73E-03 | CD36;TLR4 |
| GO:0072332 | intrinsic apoptotic signaling pathway by p53 class mediator | 1.24E-02 | MEF2C;CLDN3 |
| GO:0006310 | DNA recombination | 1.77E-02 | MEF2C;CTNNB1; BRCA1 |
| GO:0051091 | positive regulation of sequence-specific DNA binding transcription factor activity | 1.89E-02 | BRCA1;BRCA2 |
| GO:0060070 | canonical Wnt signaling pathway | 2.56E-02 | E2F1;CTNNB1;TP53 |
| GO:0000165 | MAPK cascade | 3.23E-02 | ERBB2;CD36;EZH2 |
| GO:0031325 | positive regulation of cellular metabolic process | 0.04 | DVL1;E2F1;TP53 |
| GO:0007169 | transmembrane receptor protein tyrosine kinase signaling pathway | 4.51E-02 | BRCA1;TP53 |
| GO:0048012 | hepatocyte growth factor receptor signaling pathway | 4.54E-02 | MEF2C;ERBB2;CTNNB1 |
| GO:0042982 | amyloid precursor protein metabolic process | 4.68E-02 | BRCA1 |
| GO:0034142 | toll-like receptor 4 signaling pathway | 4.94E-02 | MEF2C |

## 4 DISCUSSION

We used univariate, multivariate Cox PH ratio analysis using mRNA expression data, estimating the survival curve using product limit procedure and determining whether there is any statistically significant difference between the altered and un-altered groups using log rank test for each gene. This identified four significant genes (CD36, KLK6, MEFC2 and SCGB2A1) in univariate, three (KLK6, CD36 and E2F1) in multivariate and six genes(WFDC2, E2F1, BRCA1, KLK6, SCGB2A1 and SLPI) in combined analysis.

KLK6 gene was found significant in all three anlaysis. It is a member of kallikrein-related peptidase family and found to be frequently dysregulated in OCand responsible for modulation of tumor growth, migration, invasion, and resistance to chemotherapy (Ahmed et al. (2016)). Previously KLK6 was shown to be associated with aggressiveness of ovarian cancer (Seiz et al. (2012); Ahmed et al. (2016)). Members of kallikrein-related peptidase family were found to be significant biomarker in prostate, breast, ovary, and testis cancer (Diamandis and Yousef (2002)). Angiogenesis, regulation of cell proliferation, regulation of cell cycle process are all significantly associated with OC (Lu and Lu (2017)). KLK6 is involved in regulation of cell differentiation, tissue regeneration and hormone metabolic processes (UniProtKB) and thus is a promising candidate OC biomarker.

CD36 is clearly implicated not only in OC, but in development of other cancers (Jia et al. (2018)). From KEGG pathway analysis, this gene is found actively involved in Adipocytokine signaling pathway, PPAR signalling pathway, and AMPK signaling pathways. In other studies it found to be a significant gene affecting survival of patient of ovarian cancer (Ladanyi et al. (2018); Cuello et al. (2018)).

SCGB2A1 (Secretoglobin Family 2A Member 1) is a top differentially expressed gene in all grades of OC (Bellone et al. (2013); Fischer et al. (2014)). In addition this gene has been significantly associated with breast cancer (Zafrakas et al. (2006)) and Hypotrichosis (Tanahashi et al. (2014)).

E2F is a family of transcription factors that is recognized to regulate the expression of genes essential for a wide range of cellular functions, including cell cycle progression, DNA repair, DNA replication, differentiation, proliferation, and apoptosis. E2F1 is the most classic member of the E2F family. This gene exhibits a complex role in tumor development regulation. It was shown that E2F1 is associated with ovarian carcinoma (Zhan et al. (2016)) and a promising candidate for drug target in ovarian carcinoma. From the KEGG pathway analysis, it was found that this gene is actively involved in non-small cell lung cancer, Chronic myeloid leukemia, Cell cycle pathways. E2F1 have MicroRNAs (miRNAs) binding site which is illustrated to have significant association with cancer (Katz et al. (2015); Zhao et al. (2013)). Mutations in BRCA1 and BRCA2 is experimentally implicated in OC Sowter and Ashworth (2005). The PI3K/AKT/mTOR pathway is activated in approximately 70% of ovarian cancer cases (Gasparri et al. (2017)). From KEGG pathway analysis, BRCA1 was found to be involved in PI3K/AKT/mTOR pathway.

Previous studies have found that the MEF2C gene expression is reduced in most patients with stage III serous ovarian cance (Kim et al. (2010)). MEF2C is a transcription factor involved signalling pathways associated with OC (Estrella et al. (2014)) but is also implicated in mental retardation, autosomal Dominant 20 and arrhythmogenic right ventricular dysplasia diseases. Among its related pathways are NFAT and Cardiac Hypertrophy and Organelle biogenesis and maintenance (GeneCards). Gene Ontology (GO) annotations related to this gene include DNA binding transcription factor activity and protein heterodimerization activity (GeneCards). From KEGG pathways, we found that this gene is actively involved in Oxytocin signaling pathway inhibition of which decreases the risk of ovarian cancer in SKOV3 cells (Ji et al. (2018)), cGMP-PKG signaling pathway, Transcriptional misregulation in cancer, and MAPK signaling pathways. Previously, it was shown that the cyclic GMP (cGMP)/protein kinase G type-I (PKG-I) signaling pathway plays an important role in preventing spontaneous apoptosis as well as promoting cell proliferation in some types of cancer cells, including OC (Wong et al. (2012)).

## 5 CONCLUSIONS

In this study, survival analysis of 577 OC patients revealed that out of 26 genes chosen for their previously idedntified involvment in OC, altered transcript levels of eight of these predicted reduced OC survival. Using product limit or Kaplan-Meier analyses, we found a significant difference in survival time between altered and non-altered genes. In cases of CD36, E2F1, KLK6, SCGB2A1 and MEF2C genes, patients with these altered expression of these five genes had significantly lower survival time than patients with non-alteration of these four genes. When taking consideration of mutual effect of all selected 26 genes in multivariate analysis, CD36, E2F1 and KLK6 exhibited significant influence on survival of OC, suggesting that they may explain any survival alteration predicted by SCGB2A1 and MEF2c levels. KLK6 was notable for its high statistical significance in the Kaplan-Meier survival analysis, making this gene of particular interest, at least in this OC patient cohort. When we extended our analysis to evaluate which clinical factors and genes play dominant roles in determining survival time in OC patient, and considering interaction between clinical variables and genes, six genes including KLK6, E2F1, SCGB2A1, WFDC2, BRCA1 and SLP1 were found to be the genes having most influence on survival, i.e., their influence was independent of clinical features so would be useful to use in patient survival prediction. Again, KLK6 seems the most promising candidate for further investigation since it is significant in all three types of analysis we performed. It may be a candidate for therapeutic drug discovery itself or it may point the way to a crucial cell pathway that influences patient survival

Among the clinical variables, patient age, tumour site and cancer stage status are associated with significantly more risk of death withing 5 years in OC. In addition, we found that patients with right site tumour in ovary had more risk of death. Our approach can be used in case of other types of cancers to identify key genetic and clinical factors in patient survival.

## Abbreviations

OC: Ovarian Cancer
TCGA: The Broad Institute Cancer Genome Atlas

